# Histone H4 acetylation regulates behavioral inter-individual variability in zebrafish

**DOI:** 10.1101/100230

**Authors:** Angel-Carlos Román, Julián Vicente-Page, Alfonso Pérez-Escudero, Jose M. Carvajal-González, Pedro M. Fernández-Salguero, Gonzalo G. de Polavieja

**Author notes:** Contributed equally (Angel-Carlos Román, Julián Vicente-Page).

## Abstract

Animals can show very different behaviors even in isogenic populations, but the underlying mechanisms to generate this variability remain elusive. We found that laboratory and isogenic zebrafish (*Danio rerio*) larvae showed consistent individual behaviors when swimming freely in identical wells or in reaction to stimuli. We also found that this behavioral inter-individual variability was reduced when we impaired the histone deacetylation pathway. Individuals with high levels of histone H4 acetylation, and specifically H4K12, behaved similar to the average of the population, but those with low levels deviated from it. More precisely, we found a set of genomic regions whose histone H4 acetylation is reduced with the distance between the individual and the average population behavior. We found evidence that this modulation depends on a complex of Yin-yang 1 (YY1) and histone deacetylase 1 (HDAC1) that binds to and deacetylates these regions. These changes were not only maintained at the transcriptional level but also amplified, as most target regions were located near genes encoding transcription factors. We suggest that stochasticity in the histone deacetylation pathway participates the generation of genetic-independent behavioral inter-individual variability.

## INTRODUCTION

Classically, the phenotypic diversity of a population is considered to be generated by the genetic differences between its members and the disparity of their environmental influences (Galton, 1874). A simple prediction from this view alone would then be that isogenic populations would not show variability when the environment is constant. Nevertheless, a pioneering study showed that there was variability independent of genetic differences in some morphological traits in mice raised in identical environments (Gärtner, 1990). In recent years, similar results have been obtained for behavioral variability in mice and flies (Kain *et al*, 2012; Freund *et al*, 2013). There are several mechanisms that might contribute to this effect, including developmental noise (Waddington, 1957), maternal and paternal effects (Seong *et al*, 2011), or the different experiences the individuals obtain by interacting with the environment or other animals (Freund *et al*, 2013), among others.

Our knowledge about behavioral variability independent of genetic differences has increased substantially, but its underlying mechanisms remain unclear. Neuronal mechanisms such as neurogenesis, or serotonin signaling have been shown to be final targets of behavioral individuality (Freund *et al*, 2013; Kain *et al*, 2012), but the molecular mechanisms remain elusive. Chromatin modifications could be a promising mechanism to encode stable differences among individuals and they have been hypothesized as a potential mechanism for the generation of experience-dependent behavioral individuality (Freund *et al*, 2013). DNA methylation differences have been associated to behavioral castes in honeybees (Herb *et al*, 2012), and they are necessary and sufficient to mediate social defeat stress (Laplant *et al*, 2010). Histone acetylation is another of the main epigenetic modifications (Grunstein, 1997) and it has been shown to regulate different behaviors such as mating preference in prairie voles (Wang *et al*, 2013) or cast-mediated division of labor in ants (Simola *et al*, 2015). We thus reasoned that molecular mechanisms linked to epigenetic modifications could lead to behavioral inter-individual variability.

We used zebrafish from 5 to 8 days post fertilization (dpf) to dissect the molecular substrates of behavioral inter-individual variability. Laboratory zebrafish larvae show individuality in behavior (Pantoja *et al*, 2016) and they present some advantages such as its wide genomic information, the simplicity of its pharmacological treatments and the possibility to do large-scale behavioral analysis. Additionally, it is relevant to use a species in which we can observe directly developmental changes, as some of the mechanisms responsible for behavioral individuality are likely accumulated during development (Fraga *et al*, 2005). Here we established zebrafish larvae as a model for the analysis of inter-individual variability in free-swimming behavior. We found that in our experimental tests behavioral inter-individual variability of zebrafish larvae is independent of the genetic differences but it is correlated to histone 4 acetylation levels in a specific set of genomic sequences and regulated by a molecular complex composed by at least YY1 and HDAC1.

## RESULTS

### Behavioral inter-individual variability in larval zebrafish is stable for days

We used three steps to establish zebrafish larvae as a model to study behavioral inter-individual variability using a high-throughput setup (see **Methods and** Figure EV1A-K for the custom-built video tracking software, downloadable from www.multiwelltracker.es). We first obtained that each larvae showed differences in their spontaneous behavior, as they can be observed by simple eye inspection of trajectories (Figure 1A-B, **from 5 to 8 dpf**). This can be quantified using two parameters: overall activity (percentage of time in movement) and radial index (average relative distance from the border towards the center of the well). These two parameters were chosen because they are independent of each other (Figure 1C, *P*=0.98), while others like speed, bursting or tortuosity significantly correlated with activity (**Methods**, Figure EV1L, *P*<0.006). We performed several control experiments to show that the observed inter-individual behavioral variability was not affected by potential artifacts in the setup (**Methods**, Figure EV2A-C). Also, we tested that these behavioral parameters describe inter-individual variability in response to stimuli like light flashes, mechanical perturbation or at a novel tank (**Methods**, Figure EV2D).

**Figure 1.**
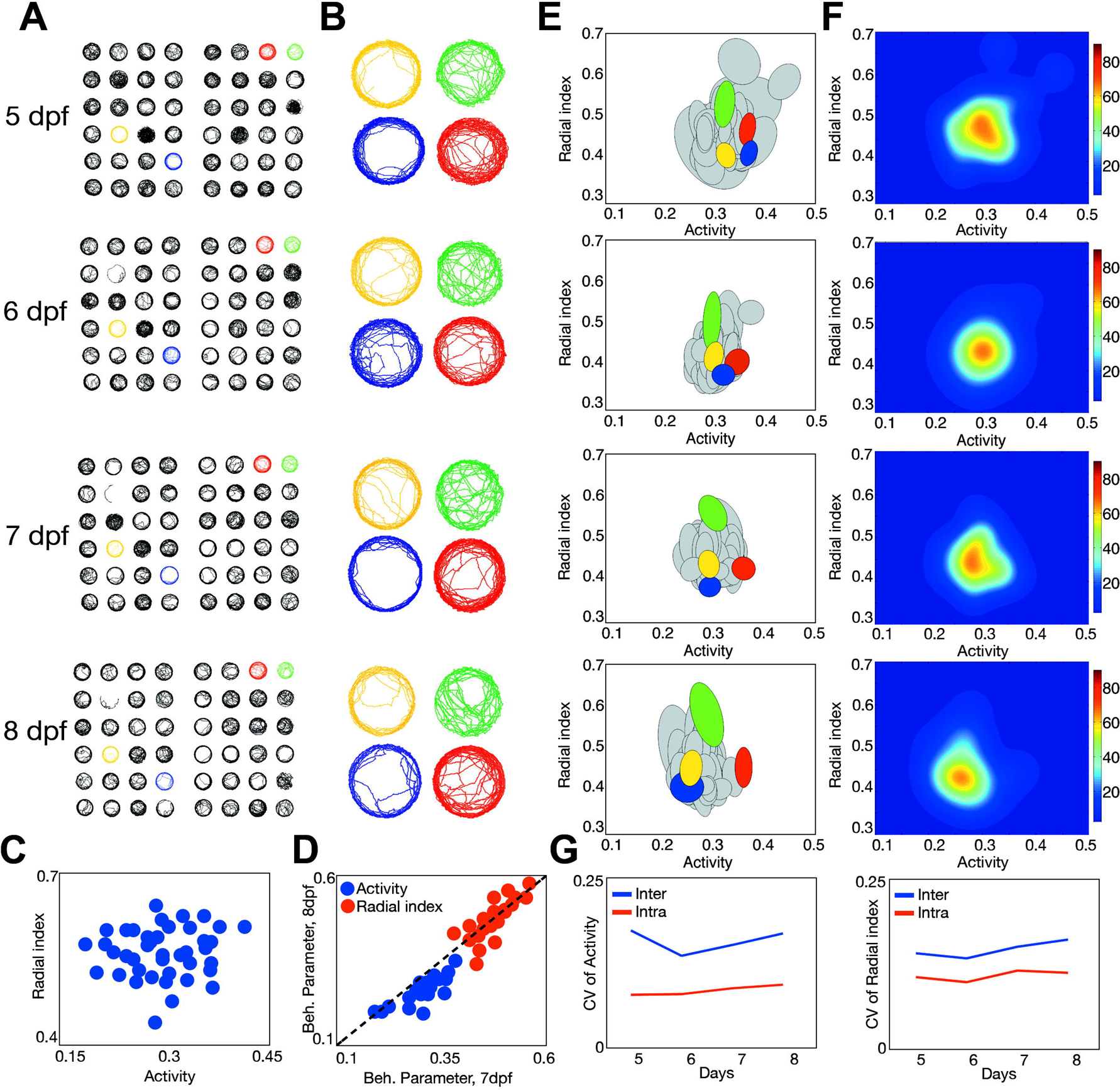
Behavioral inter-individual variability in a population of 48 larval zebrafish. **(A)** Example 20-minute trajectories for the same larval group recorded at 5-8 dpf. **(B)** Trajectories from four specific larvae zoomed from A. **(C)** Radial index *vs*. activity at 7 dpf of the same group. **(D)** Correlation of activity (blue) and radial index (red) between 7 dpf and 8 dpf for the same group. **(E)** Population variability in activity and radial index of the same group at 5-8 dpf. Each ellipse represents the behavioral intra-individual variability for each single fish as described in **Methods**. Colors as in **A and B**. **(F)** Probability density of finding an individual with a given mean activity and radial index at 5-8 dpf. **(G)** Median of intra-individual variability (red) and inter-individual variability (blue) for activity (left) and radial index (right) during the time course of the experiments.

In a second step, we showed that individual differences were robust along several days (Figure 1D, R=0.69 and R=0.58, *P*<0.001 for linear correlation of activity and radial index, respectively, 7 vs. 8 dpf; Figure EV2E, R=0.48 and R=0.41, *P*<0.01, 5 vs. 6 dpf). Finally, the third step consisted in showing that inter-individual variability is larger than intra-individual variability. This can be directly seen in the two-dimensional phenotypic space defined by activity and radial index, by noticing that the area covered by the behavior of one individual is smaller than the area of the whole population (Figure 1E, see **Methods**). Measuring variability by the Coefficient of Variation (CV), we found this difference to be significant (**Methods**, Figure EV2F-I, *P*<0.001 in all cases). Intra-individual and inter-individual variability levels remained stable from 5 to 8 dpf (Figure 1G, *P*<0.01 in all comparisons).

The degree of inter-individual variability can be visualized using the probability density of finding an individual in a population with a given mean activity and radial index (**Methods**, Figure 1F). It can be quantified using *generalized variance* (Wilks, 1932), a single parameter we used to compare two populations that summarizes this two-dimensional variability (**Methods**; **Table 1** for summary of all results; other variability measures in **Table 2**).

### Sources of behavioral inter-individual variability in zebrafish

Our setup allowed us to perform high-throughput tests to study the possible origins of behavioral inter-individual variability, which could in principle depend on environmental manipulations and the genetic differences across the population. Our experiments minimized environmental influences by isolating eggs in plates at pharyngula stage (24 hpf) and by keeping them at a controlled temperature (27-28ºC). Manual changes in water (24 hours before the experiment) or feeding did not affect inter-individual variability (Figure 2A, *P*=0.42 and Figure 2B, *P*=0.38, respectively).

**Figure 2.**
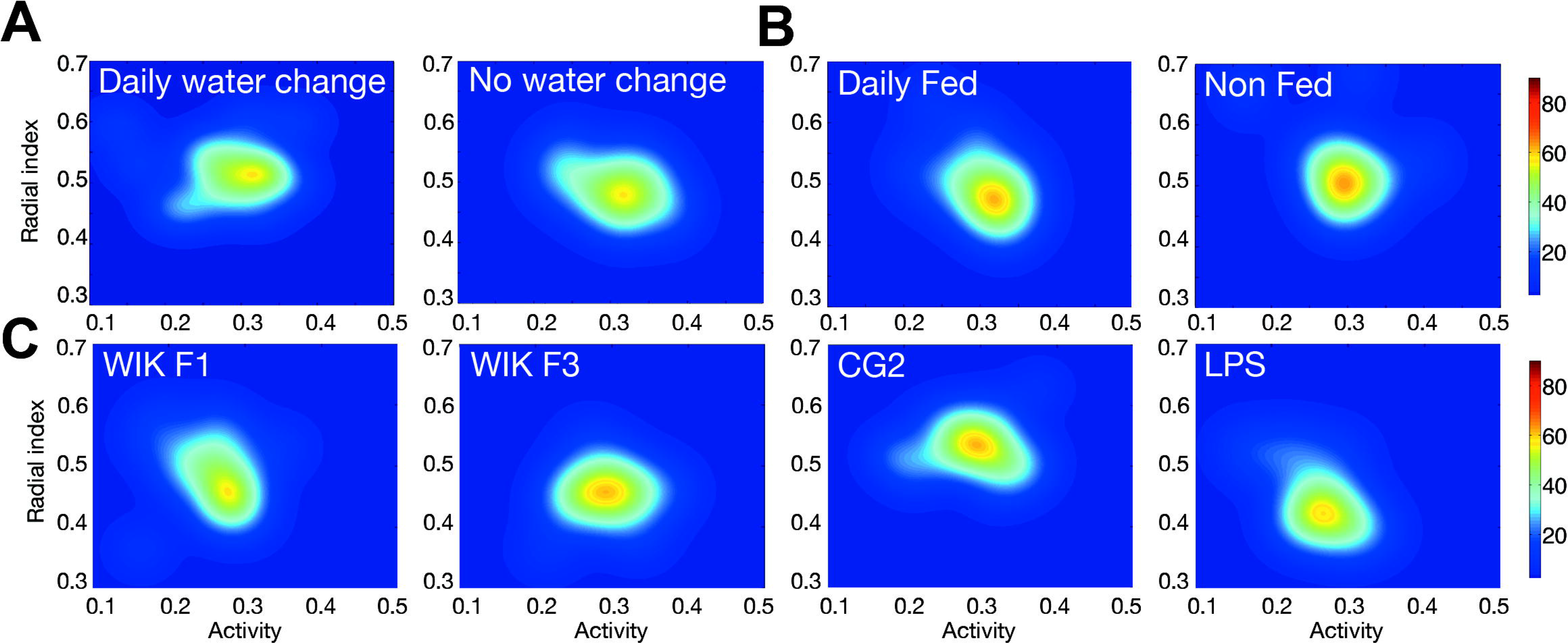
Impact of environmental changes and genetic background on behavioral inter-individual variability. **(A)** Probability density of finding an individual with a given mean activity and radial index for additional larval groups (24 individuals) with and without daily water changes, at 7 dpf. **(B)** Same as **A**, but for additional daily fed (control case) and non-fed animals throughout the experiment **(C)** Same as **A** for additional groups with different genetic backgrounds: WIK F1 (three inbreeding cycles), WIK F3 (five inbreeding cycles), CG2 (gymnozygotic fish clones) and LPS (outbred parents).

We also found that behavioral variability of a population did not depend on the genetic variability of its individuals. Our control laboratory WIK zebrafish population (F1) resulted from a single batch of eggs retrieved from two adults with at least three cycles of inbreeding. We obtained the same behavioral inter-individual variability after two more inbreeding cycles (WIK F3, Figure 2C, *P*=0.33) and in an isogenic population (Mizgirev & Revskoy, 2010) (CG2, Figure 2C, *P*=0.44). Also, we did not find changes in the behavioral inter-individual variability using groups of siblings from genetically diverse outbred parents (LPS line, Figure 2C, *P*=0.38).

### Changes in histone acetylation pathway alter behavioral inter-individual variability

The absence of effects from genetic variability prompted us to test whether behavioral inter-individual variability could be modified by different epigenetic factors. To test the contribution of DNA methylation we used 5-azacytidine (AZA), an inhibitor of DNA-methyltransferases (Christman, 2002). We found that AZA added to the water did not alter the behavioral inter-individual variability of a population (15 mM AZA, Figure 3A, *P*=0.44) even if it reduced 3-methyl DNA in larval zebrafish (Figure EV3A). We then studied the role of histone deacetylation, a reversible molecular process in which an acetyl functional group is removed from specific residues of Histone H3 and H4 (Grunstein, 1997). This system is regulated by a group of enzymes called Histone Deacetylases (HDACs) that can be divided into three classes based on their sequence homology. Class I HDACs (HDAC1, HDAC3 and HDAC8 in the case of the zebrafish) are strictly localized in the cell nucleus, and they are normally ubiquitously expressed, while Class II HDACs shuttle from cytoplasm to nucleus, and each protein is specifically expressed in a few tissues. Class III enzymes are different from class I and II from a phylogenetic point of view, they are NAD^+^-dependent deacetylases, and known as sirtuins (Ruijter *et al*, 2003). To test the effect of HDACs on behavioral inter-individual variability, we first used sodium butyrate (NaBu, a class I HDAC inhibitor) at the standard concentration of 2 mM (Heruth *et al*, 1993), and we confirmed that it increases the level of total acetyl-histone H4 in larval zebrafish (Figure EV3B, *P*<0.01, NaBu). We found that this treatment reduced the behavioral variability of a WIK F3 sibling population after 24 hours (2 mM NaBu, Figure 3B, **top right**) compared to control PBS-treated larvae (PBS, Figure 3B, **top left**, *P*<0.001). Note that this treatment only altered variability and not the mean of the population parameters (*P*=0.63). When we removed the NaBu from the water, behavioral variability was recovered after additional 24h (Figure EV3C, *P*=0.71). Similarly to the behavior, the total levels of acetyl-histone H4 increased with the treatment and were recovered 24 hours after washing the medium (Figure EV3B, *P*=0.42, NaBu/PBS). Another HDAC inhibitor (against class I and class II HDACs) like Trichostatin A (0.1 μM TSA, Figure 3B, **bottom left**) had a similar behavioral effect as NaBu, reducing the behavioral variability of the population (*P*=0.02) and increasing histone H4 acetylation (Figure EV3B, *P*<0.01, TSA). In contrast, a inhibitor of class III HDACs like cambinol (0.2 μM Cambinol, Figure 3B, **bottom right)** did not alter the behavioral variability of the population (*P*=0.71), even when cambinol treatment increased the acetylation levels of histone H4 (Figure EV3B, *P*<0.01, Cmb). We confirmed the specificity of the effects on class I histone deacetylases by studying transgenic larvae as an alternative to the use of drugs. We found that two different heterozygotic mutant populations of the class I histone deacetylase *hdac1* (*hdac1* +/−), *sa436* (Noël *et al*, 2008) and *hi1618* (Golling *et al*, 2002), showed a reduced behavioral inter-individual variability compared to their AB controls and an increase in histone H4 acetylation, mirroring the results obtained with the drugs (Figure 3C **top**, and **bottom left**, *P*=0.008 for *sa436* and *P*=0.006 for *hi1618*; Figure EV3B, *P*<0.01, hdac1 +/−). Addition of 2 mM NaBu to the *hdac1* +/− *hi1618* population did not change the behavioral inter-individual variability of the larvae (Figure 3C **bottom right**, *P*=0.23), confirming that the drug treatment is affecting this variability through HDAC1. Our results suggest that the histone deacetylation pathway modulates the behavior of zebrafish larvae without affecting its average behavior.

**Figure 3.**
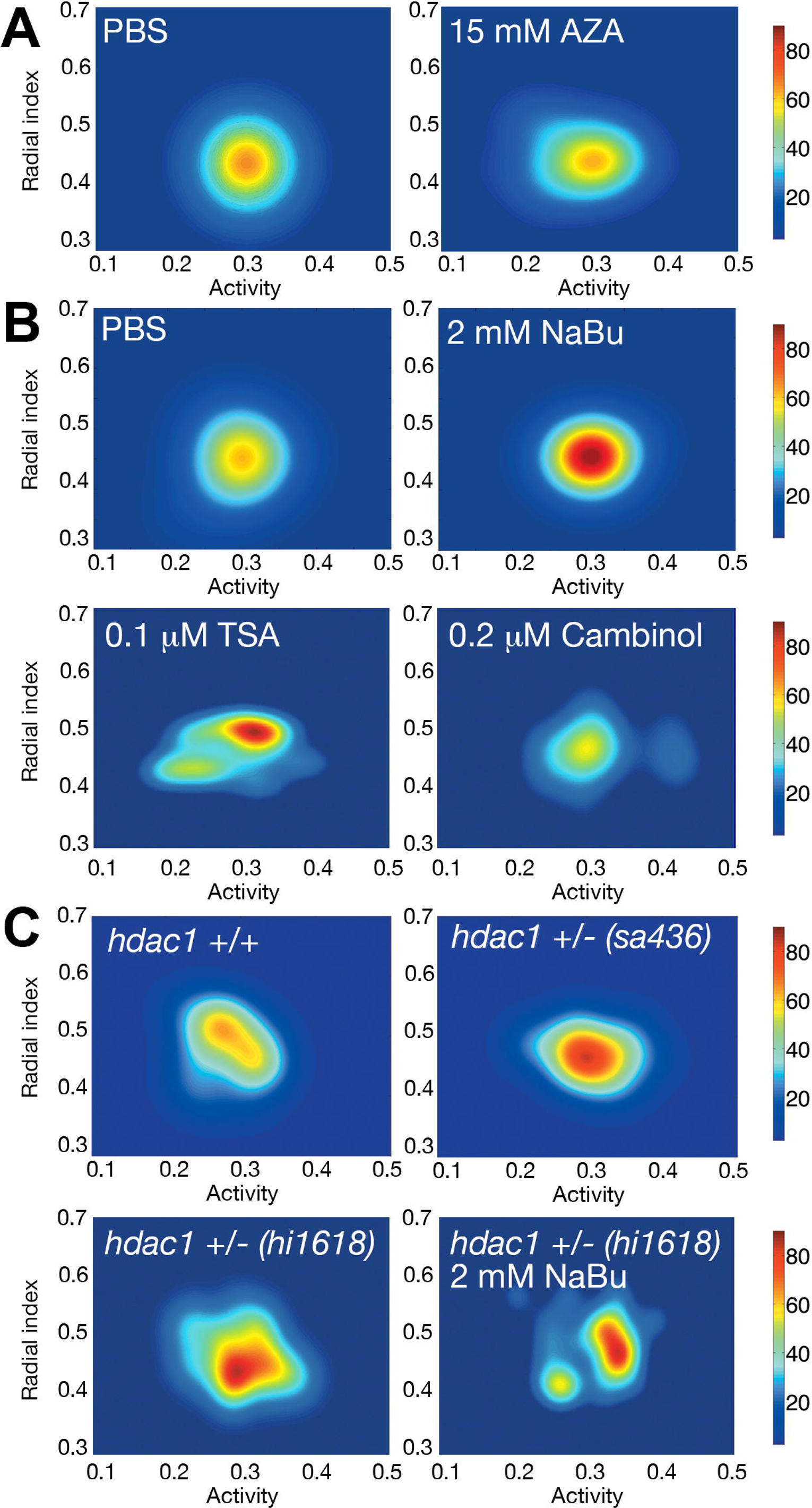
Epigenetic modulation of behavioral inter-individual variability. **(A)** Probability density map for 24 fish treated with a PBS solution as control and 15 mM AZA for 24 hours. (**B**) The same for PBS (top left), 2mM NaBu (top right), 0.1 μm Trichostatin A (bottom left) and 0.2 μm Cambinol (bottom right). **(C)** Probability density map for *hdac1* +/+ (top left), *hdac1* +/− (sa436 mutant, top right), *hdac1* +/− (hi1618 mutant, bottom left) and *hdac1* +/− (hi1618 mutant) with 2 mM NaBu for 24 hours (bottom right) larvae.

### Histone H4 acetylation levels correlate with behavioral distance to average behavior

We have shown that the degree of behavioral inter-individual variability of a population depends on its average acetylation levels. Since an increment in the global histone acetylation decreased this variability without changing the average behavior, we reasoned that the individuals with higher mean acetylation should be placed near the average population behavior in the phenotypic space. To test this hypothesis, we performed an experiment with 90 zebrafish individuals to obtain their histone H4 acetylation state depending on their distance to the average behavior of the population. As we needed at least five larvae in order to get enough tissue for the experiment, we pooled 5 larvae with very similar behavior and measured their acetylation state using ELISA kits that allow the quantification of histone H4 acetylation and total histone content. We found that pools of larvae whose behavior was placed near the average of the population had higher mean histone H4 acetylation values (Figure 4A, **left**, *P*=0.007). To quantify the dependence between the average histone H4 acetylation and the position in the phenotypic space of the samples, we first defined a coordinate system (centered on the average behavior of the population) and then obtained two magnitudes for each pool of fish: their average distance to the center (*r*) and their average angle with the horizontal axis (*θ*) (Figure 4A, **right**). We found that the histone H4 acetylation levels of the pools highly correlated with their phenotypic distance *r* to the average, while we found no correlation with their angular position *θ* (see Figure 4B, blue dots, *P*<0.001 and *P*=0.53, respectively). We found similar correlations when we analyzed the distance to the mean of each behavioral parameter separately (Figure EV4A, *P*<0.001 for both activity and radial index).

**Figure 4.**
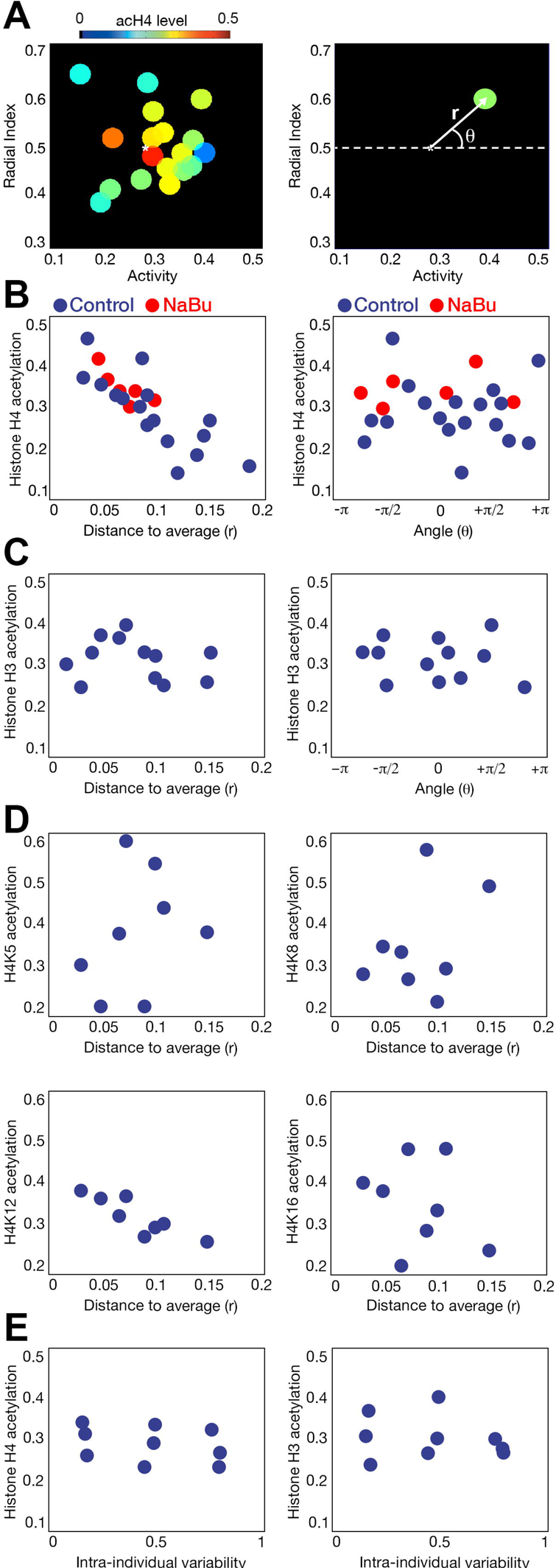
Relation between histone acetylation levels and behavior. **(A)** Average histone H4 acetylation levels of fish depending on their behavior using 90 fish as the initial population and clustering them into groups (left). Schematic representation of the two parameters used to analyze the dependence between histone H4 acetylation and behavior (right). **(B)** Relation between the values of histone H4 acetylation and the two parameters of the coordinate system centered on the average behavior of the population: the distance to the average (left) and the angle with the horizontal axis (right). Blue dots are control larvae while red dots indicate Nabu-treated animals. (**C**) Same as B, but for histone H3 acetylation. (**D**) Relation between the values of histone H4K5 (top left), H4K8 (top right)m H4K12 (bottom left) and H4K16 (bottom right) acetylation and the distance to the average behavior of the population. (**E**) Relation between the values of histone H4 (left) and H3 (right) acetylation and clusters of larvae with similar intra-individual behavioral variability.

So far, we have shown that individuals with higher histone H4 acetylation levels display a behavior similar to the average of the population, while the variability of the population behavior increases at lower histone H4 acetylation levels. This is consistent with our previous experiments that reduced the behavioral variability of a population by increasing its acetylation levels using HDACi or *hdac1* +/− mutant animals. In fact, when we used fish treated with NaBu to perform the same acetyl-H4 quantification, we found that their histone H4 acetylation is at a similar level to the non-treated individuals with highest histone H4 acetylation (Figure 4B, red dots, *P*=0.24). This shows that the NaBu-treated animals present histone H4 acetylation levels within the physiological range of the animals, consistent with NaBu having the global effect of increasing the histone H4 acetylation levels of the population by bringing them close to the animals with highest acetylation. Then we analyzed if the effect observed for histone H4 acetylation was also present in histone H3. Interestingly, histone H3 acetylation levels were correlated neither to *r* nor *θ* (Figure 4C, *P*=0.61, and *P*=0.64, respectively), This result made us focus on the specific marks for histone H4 acetylation. We found that H4K12 acetylation levels correlated with *r*, while other marks were not found to be affected by the behavioral position of the samples (Figure 4D and EV4B, *P*=0.005 for H4K12, *P*>0.27 for the rest of marks). Our approach consisting in pooling larvae with a similar behavior is consistent, as we observed that different pools obtained from the same behavioral position maintain very similar acetyl-H4 and acetyl-H3 levels (Figure EV4C, **left**), and pools located very near one another in the behavioral space maintain very similar histone H4 acetylation levels compared to the rest of clusters (Figure EV4C, **right**, *P*=0.56).

Finally, we studied if intra-individual variability could be also linked to histone acetylation. We pooled samples with similar intra-individual variability and we quantified histone H4 and H3 acetylation (Figure 4E). We did not find a correlation between acetylation levels and behavioral intra-individual variability (*P*=0.49 and *P*=0.58, for histone H4 and H3, respectively).

### Genomic regions linking histone H4 acetylation and behavioral inter-individual variability

Our results link histone H4 acetylation level of the individuals to their behavior. However, fish with similar (low) histone H4 acetylation levels also can show very different behaviors, so there must be other factors contributing to behavioral inter-individual variability. We hypothesized that these factors could be the acetylation differences in specific genomic regions associated to behavior. To explore this possibility, we compared the histone H4 acetylation levels between two groups of zebrafish, one with high and the other with low behavioral variability. For the first population, we used four samples of five pooled sibling fish with similar behavior (see **Methods** for details). For the second population, we used four samples of five sibling fish treated with NaBu with similar behavior. We then retrieved the acetyl-H4 epigenomic profiles of the samples in each group using chIP-seq and calculated the peaks for each sample with the MACS algorithms. Then, we computed the histone H4 acetylation differences for each peak between control and NaBu groups using standard techniques adopted from gene expression analysis as EdgeR (see Figure 5A and **Methods** for details). Finally, the candidate peaks with significantly higher histone H4 acetylation in the NaBu group (*P*<0.01) were filtered to remove the regions whose acetyl-H4 variability were not associated to behavior. In order to do this, four samples of five randomly pooled fish were subjected again to acetyl-Histone H4 chIP-seq, and the acetylation levels for the previously candidate peaks were obtained. As the variability in this group was not associated to behavior because the fish were selected randomly, we rejected candidate peaks in which their histone H4 acetylation variability were equal or higher than in the control group (Figure 5A). From this procedure we obtained a final set of 729 regions in which sodium butyrate increased histone H4 acetylation potentially responsible for the behavioral inter-individual variability differences observed after alteration of the HDAC pathway (**Table 3**).

**Figure 5.**
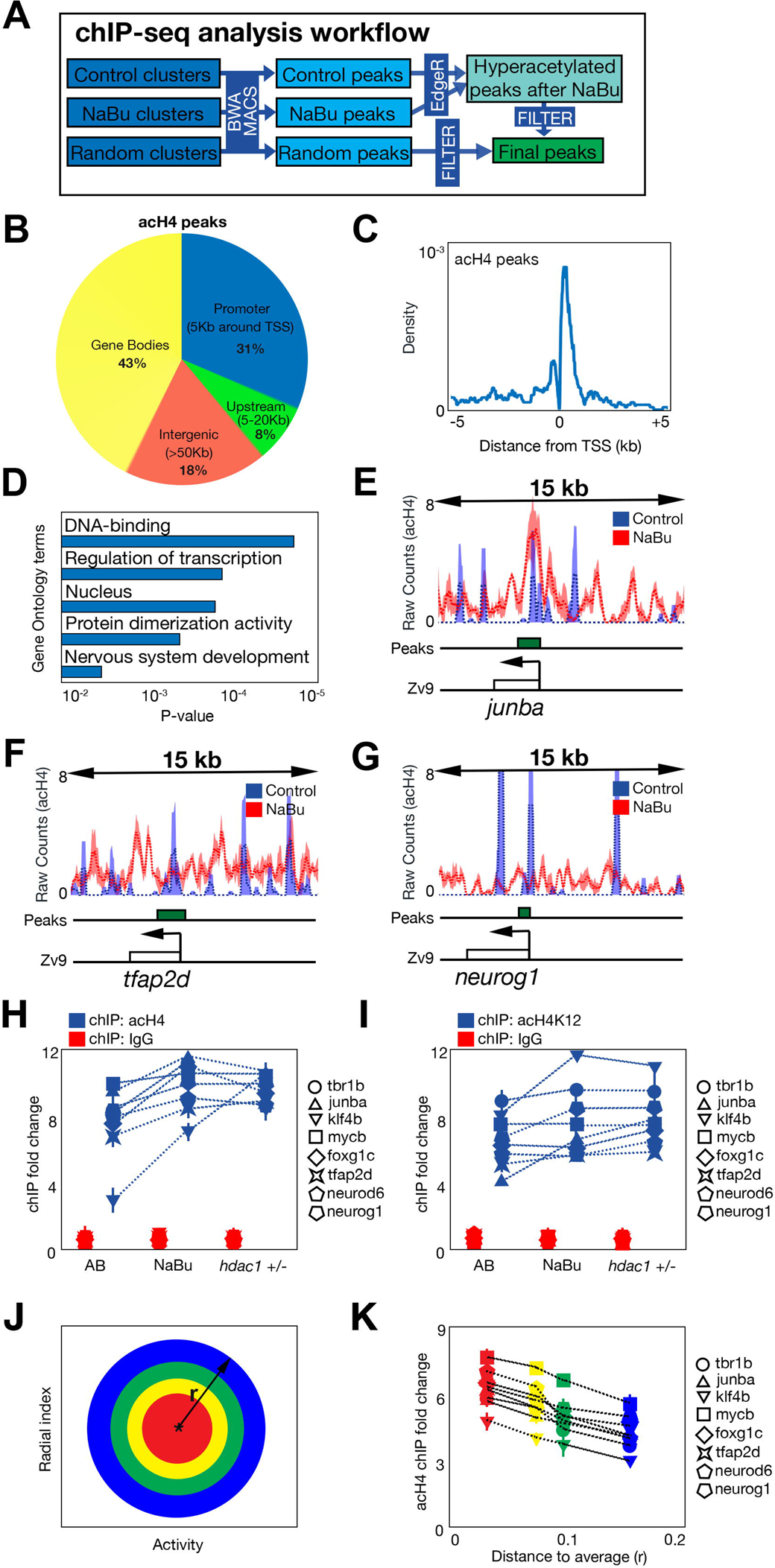
Histone 4 acetylation regions related to behavioral inter-individual variability. **(A)** Workflow of the analytical steps using histone H4 acetylation chIP-seq. **(B)** Classification of the acetyl-H4 peaks obtained depending on their relative position to the transcription start site (TSS) of the nearest gene. (**C**) Histogram representing the relative positions of the acetyl-H4 peaks located around the TSS of the nearest gene. (**D**) Enriched gene ontology terms of the acetyl-H4 peaks. (**E**) Snapshot of the raw reads results obtained in the acetyl-H4 chIP-seq in the region around *junba* (marked with a box and an arrow showing its TSS). Blue dotted line indicates the mean reads in the four control samples, while red dotted line indicates the mean reads in the four NaBu-treated samples. Blue and red areas show the 25%-75% range of the reads in control and NaBu samples, respectively. Green box indicates the detected peak by the algorithm. (**F**) Same as E, but for tfap2d gene. (**G**) Same as E, but for neurog1 gene. (**H**) Histone H4 acetylation levels quantified in conventional chIP as the fold change compared to non-bound fraction in eight selected regions in control, NaBu-treated and *hdac1* +/− larvae. Blue figures represent the acetyl-H4 chIP, while red figures show a control chIP using only IgG. Bars represent standard deviation using three replicates. A legend in the right indicates the names of the regions. (**I**) Same as **H**, but using H4K12 acetylation chIPs. (**J**) Diagram representing cluster selection in **K**. Each color marks the behavioral area from which we will retrieve larvae for comparison. This area depends on the distance to the average behavior of the population (*r*). (**K**) Same as **H**, but comparing the acetyl-H4 differences in the different clusters represented in **J**.

We studied the relative genomic positions of these peaks (Figure 5B). We found that these peaks, that were hyperacetylated after NaBu, were enriched in promoter regions (+/− 5 Kb around the Transcription Start Site (TSS)) and gene bodies (31% and 43%, respectively). Upstream regions (from 5 Kb to 20Kb of the TSS) accounted for 8% of the total peaks, while intergenic peaks were 18%. This distribution is not very different to others obtained for acetyl-histone marks in other conditions (Karmodiya *et al*, 2012). We then studied the subset of peaks located near the TSS, as the histone H4 acetylation changes detected in these regions could be associated to differences in the expression of near genes. We observed that these peaks were located very near to the TSS, but excluded from the exact TSS (Figure 5C), a typical acetyl-H4 effect seen in a previous work (Karmodiya *et al*, 2012). We then characterized this set of regions located near TSS regarding the Gene Ontology (GO) terms associated with their neighboring genes (Figure 5D). We found five enriched (*P*<0.01) terms, the highest related to transcription factor activity, so we selected eight of the candidate peaks that were located near a transcription factor (*tbr1b, junba, klf4b, mycb, foxg1c, tpa2d, neurod6, neurog1*) to assess their role in the control of behavioral inter-individual variability (see Figure 5E-G for chIP-seq snapshots of the normalized levels of acetyl-histone H4 around these regions).

We analyzed the levels of histone H4 acetylation in these eight regions by conventional chIP at three different conditions: AB populations, NaBu-treated animals, and *hdac1* +/− populations (Figure 5H). We observed that the acetyl-H4 content in these regions was not only increased after NaBu treatment (*P*<0.01 for all the regions), as predicted by the chIP-seq results, but also in *hdac1* +/− populations (*P*<0.01 for all the regions). Acetylation of the specific H4K12 mark in these regions did not reflect exactly the same results of global acetyl-H4 (Figure 5I), as five regions increased their H4K12 acetylation in either NaBu-treated or *hdac1* +/− populations (*tfap2d*, *junba*, *klf4b*, *neurog1* and *tbr1b*, *P*<0.03), while two of them only increased H4K12 acetylation in *hdac1* +/− larvae (*foxg1c*, *neurod6*, *P*<0.02) and another one did not respond at all (*mycb*). These results suggest that the effect of NaBu was mediated by the inhibition of HDAC1 deacetylation of H4K12 residues on specific regions of the genome. At this point, we wondered if the histone H4 acetylation levels in these eight regions reflected the effect observed for global histone H4 acetylation and behavior shown in Figure 4B: an inverse correlation between acetylation and the distance to the average behavior of the population. We prepared four pooled samples (ten larvae per sample) with different behavioral distance to the average of the population (Figure 5J), and we quantified the acetyl-H4 content within the eight selected regions (Figure 5K). We found that the histone H4 acetylation levels in these regions decreased with the distance to the average behavior (*P*<0.01 for all regions), which suggested a role for these regions in the behavioral phenotypes previously observed.

### A complex composed by YY1/HDAC1 deacetylates histone H4 in target regions

To find whether these regions could have a causal action in behavioral inter-individual variability, we decided to affect them by impairing DNA-interacting proteins that significantly bind near these regions. We found several DNA motifs that were enriched (*P*<0.0001) near the candidate regions, Yin-Yang 1 (35% of the total sequences), RUNX1 (22%), NFY (11%) binding sites and even an unknown sequence (5%) (Figure 6A). Ying-Yan 1 (YY1) is a transcription factor that can activate or repress the same target gene depending on recruited co-factors (Shi *et al*, 1991), being HDAC1 (Yao *et al*, 2001) one of its main partners. We then studied if YY1 might be implicated in the inter-individual variability by testing the behavior of a heterozygotic mutant *yy1a* (*yy1* +/−) population (Figure 6B). We found that this alteration decreased the behavioral inter-individual variability (*P*=0.003) compared to wild-type counterparts. This result suggested that YY1 is necessary for the maintenance of this variability. As YY1 binding sites are present near the candidate regions, we quantified the differences in histone H4 acetylation that occurred near the eight previously selected regions in the *yy1*+/− population. We found that, not only global histone H4 acetylation but also specific H4K12 acetylation were increased in the mutant background (Figure 6C, *P*<0.01 for all the regions except acetyl-H4K12 levels in *tbr1b*, *P*=0.23), similar to the results obtained in NaBu-treated and *hdac1* +/− animals.

**Figure 6.**
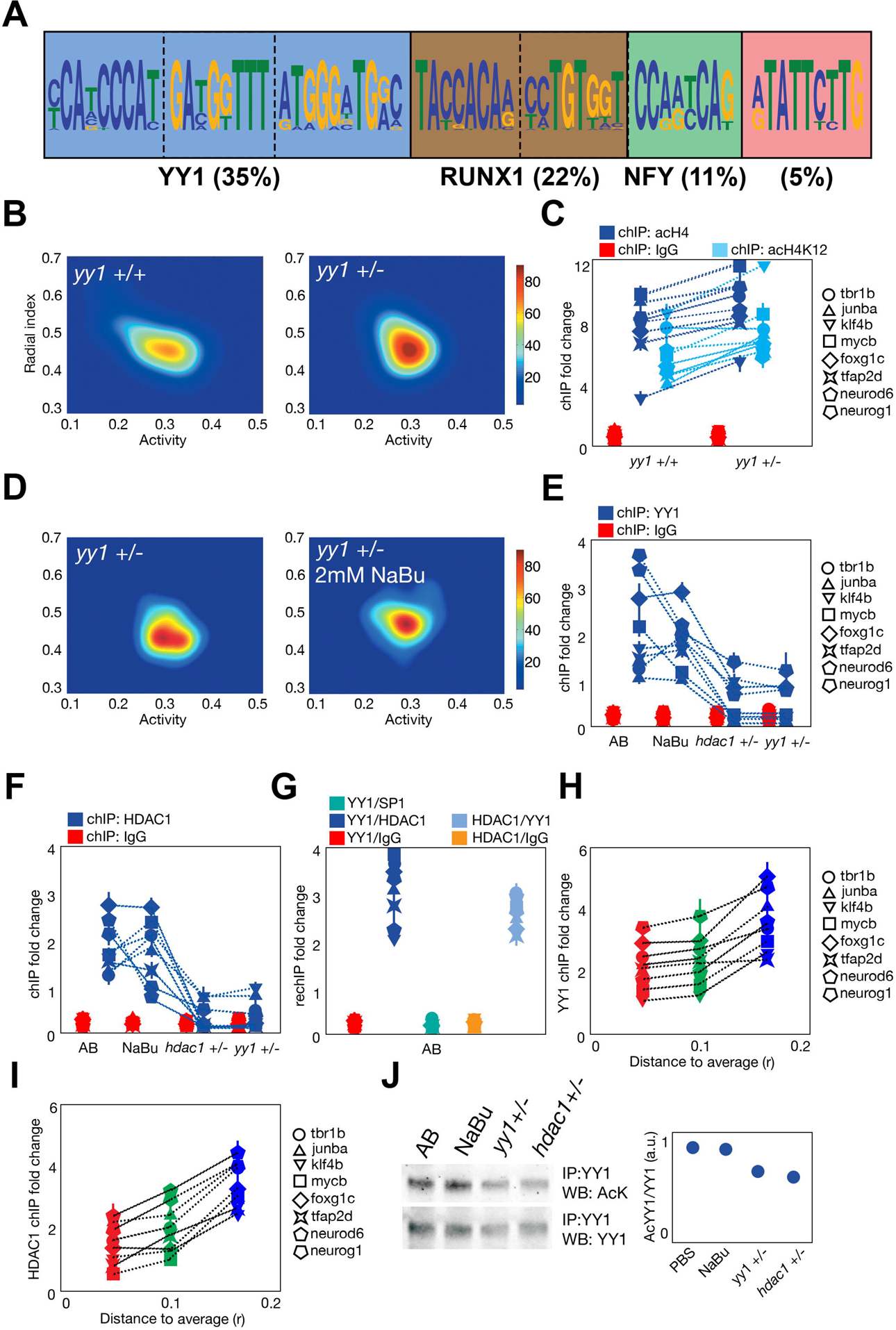
YY1 and HDAC1 role in inter-individual behavioral variability and acetyl-H4 changes. **(A)** Diagram representing the most represented motifs found in the acetyl-H4 peaks located near TSS. The predicted transcription factors that can bind to these sites are also indicated if found, with the percentage of the total peaks that present at least one of these motifs. **(B)** Probability density map for the behavior of 24 *yy1* +/+ (left) and *yy1* +/− (right) fish. (**C**) Acetyl-H4 (dark blue), acetyl-H4K12 (light blue) and additional control IgG (red) levels found in *yy1* +/+ and *yy1* +/− larvae, as detected by fold change compared to an unbound fraction in eight selected regions that are showed in the legend on the right. (**D**) The same as **B**, but for *yy1* +/− and *yy1* +/− treated with 2 mM NaBu for 24 hours. (**E**) YY1 binding (blue dots) and additional IgG presence (red dots) to the eight selected regions quantified by fold change compared to the unbound fraction in eight selected regions, in AB (control) larvae, NaBu-treated, *hdac1* +/− and *yy1* +/− fish. (**F**) Same as **E**, but for HDAC1 binding. (**G**) RechIP fold change in control AB larvae. The order of the two consecutive chIPs are noted in the names of the conditions. (**H**) YY1 binding in eight selected regions to clusters of larvae with different distances *r* to the average behavior of the population. (**I**) Same as H, but for HDAC1 binding. (**J**) YY1 acetylation in control AB larvae, NaBu-treated, *hdac1* +/− and *yy1* +/− larvae. YY1-immunoprecipitated extracts were subjected to western blot analysis using an acetylated-lysine antibody (left, top) or YY1 antibody (left, bottom). Quantification of the acetyl-YY1/YY1 ratio is shown on the right.

These results prompted us to study whether histone deacetylation and YY1 share the same pathway. First we tested the behavioral inter-individual variability of NaBu-treated *yy1* +/− animals, and a double heterozygotic *hdac1*+/− *yy1*+/− population (Figure 6D). We found that NaBu treatment did not further decrease the behavioral inter-individual variability of the *yy1* +/− fish (*P*=0.54), while the double mutant was lethal to the animals. Afterwards, we analyzed the recruitment of YY1 to the selected regions in wild-type, NaBu-treated, *yy1* +/− and *hdac1* +/− populations by chIP (Figure 6E). YY1 binds to these regions in control conditions and the treatment with NaBu did not alter this recruitment (*P*>0.05 for all the regions). Interestingly, not only in the *yy1* +/− but also in the *hdac1* +/− populations, the binding of YY1 to the regions was decreased (*P*<0.02 for all the regions), suggesting that HDAC1 presence but not its activity was necessary for the recruitment of YY1 to the candidate regions. We also performed the same experiments but on HDAC1 recruitment (Figure 6F), and it showed that the binding of the enzyme correlated to YY1 binding (*P*>0.05 for AB vs NaBu, *P*<0.01 for AB vs *yy1* +/− and AB vs *hdac1* +/−). Thus, these results not only point to a common pathway between YY1 and HDAC1, but also suggest their participation in the same regulatory complex in the candidate regions. Then we used double consecutive chIP (rechIP) to confirm our hypothesis (Figure 6G), showing that YY1 and HDAC1 were bound together to the eight regions (*P*<0.01 for all the regions). Finally, we performed the same experiment shown in Figure 5J-K, but looking at HDAC1 and YY1 recruitment in the target regions depending on the distance to the average behavior. We found that both HDAC1 and YY1 binding increases with the distance to the average behavior, consistent with the deacetylation process that occurs depending on this distance (Figure 6H-I, *P*<0.04 for all the regions). As YY1 itself can be dynamically acetylated in a process in which participates HDAC1 (Yao *et al*, 2001), we analyzed YY1 acetylation by co-immunoprecipitation in different conditions (Figure 6J and EV5). We found that YY1 is acetylated in basal conditions and a treatment with NaBu does not seem to affect this acetylation (*P*=0.51), while both in *hdac1* +/− and *yy1* +/− populations this acetylation is decreased (*P*<0.01). As YY1 acetylation does not correlate with behavioral inter-individual variability or the chromatin changes previously observed, we cannot confirm a role for this modification in the phenotype. Nevertheless, we cannot exclude its participation in the dynamics of the recruitment or the activity of the YY1/HDAC1 complex.

### Gene expression is changed in the set of regions with alterations in histone H4 acetylation

So far we have found a relationship between a YY1/HDAC1 complex, histone H4 acetylation changes and the larval behavioral inter-individual variability. Still, a mechanistic explanation is needed to describe how these chromatin alterations could lead to an altered behavior. One possible justification is that the transcriptional changes in the genes located near the candidate regions could lead to functional differences across individuals and finally to an altered behavior. To test this, we used RNA-seq to analyze if the histone H4 acetylation changes observed in chIP-seq were maintained at the gene expression level. We compared gene expression profiles retrieved for control zebrafish and NaBu-treated animals using standard techniques as TMM normalization of the read counts followed by the Noiseq algorithm to obtain a list of significantly altered genes (Tarazona *et al*, 2015) (Figure 7A, *P*<0.01). There was a significant enrichment in the overlapping between the over-expressed genes and the set of candidate peaks previously obtained in acetyl-H4 chIP-seq (25%, Figure 7B *P*=0.003).

**Figure 7.**
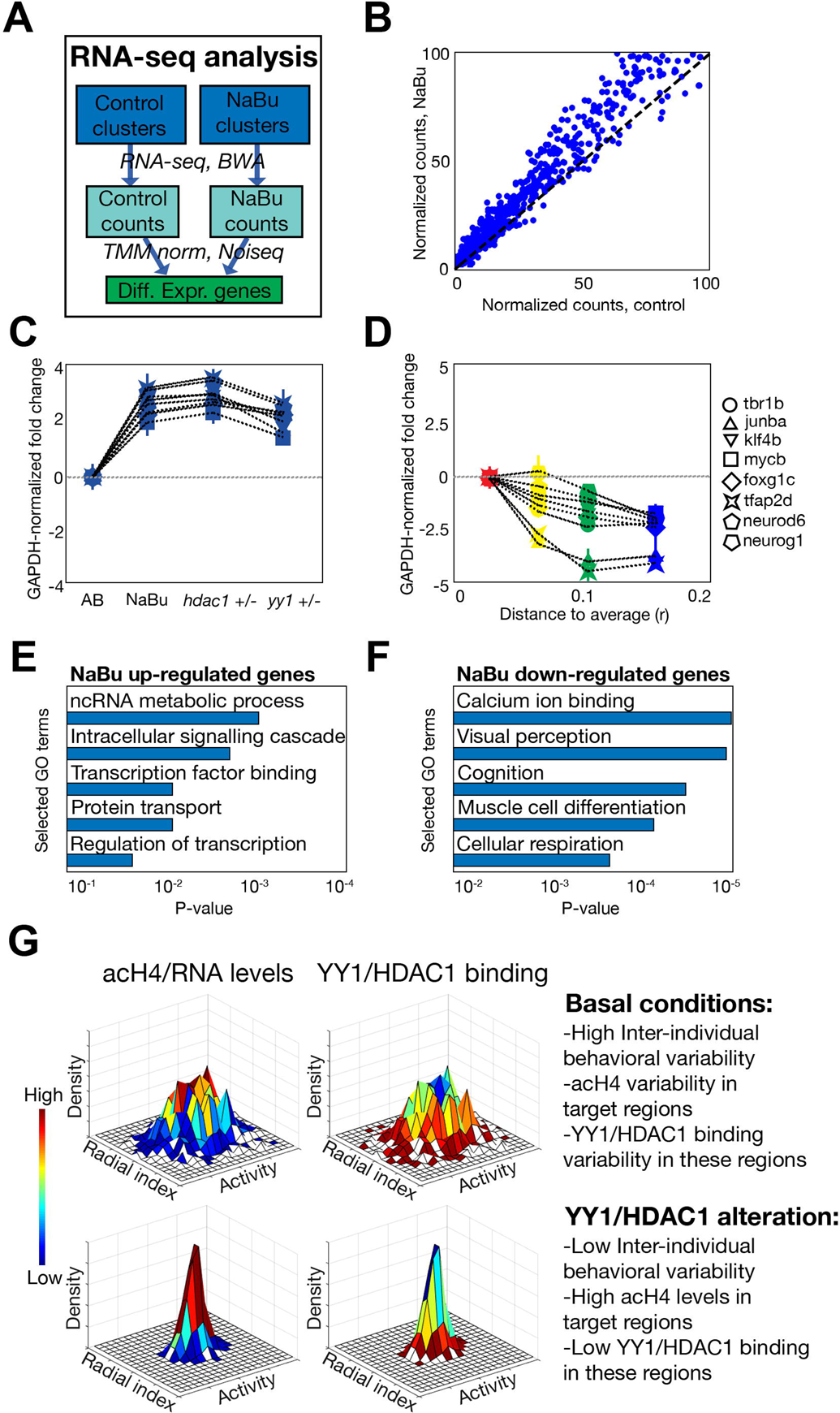
Conservation between epigenomic and transcriptomic results. **(A)** Schematic workflow of the RNA-seq analysis of control and NaBu-treated samples. (**B**) Normalized expression of the genes located nearest to the acetyl-H4 peaks detected in the chIP-seq experiment. X-axis represents control samples and Y-axis represents NaBu-treated samples. (**C**) Gene expression fold change differences in eight selected genes in AB larvae, NaBu-treated, *hdac1* +/− and *yy1* +/− larvae. Normalization was made by subtracting the values obtained for the *gapdh* gene in each sample, and bars mark standard deviation obtained in three replicates. (**D**) Same as **C**, but comparing samples with different distance r to the average behavior of the population. (**E**) Enriched Gene Ontology selected terms found in the set of genes over-expressed after NaBu treatment. (**F**) The same as E, but for genes down-regulated after NaBu treatment. (**G**) Schematic model of the results obtained in the manuscript. In control conditions, heterogeneous populations as classified by their activity and radial index are obtained. After alteration of the YY1/HDAC1 pathway, more homogeneous populations are observed, while the acetyl-H4 content and the YY1/HDAC1 presence in a set of genomic regions are anti-correlated, being these epigenetic changes transferred to the gene expression level.

We then verified the results obtained in the RNA-seq experiment by quantifying the expression of the genes located near the eight previously selected regions. As predicted by the RNA-seq and in parallel to histone H4 acetylation, gene expression was increased not only in NaBu-treated, but also in *yy1* +/− and *hdac1* +/− animals (Figure 7C, *P*<0.01 for all the genes). Moreover, gene expression decreased with the distance to the average behavior, similar to the effect that takes place at the histone H4 acetylation level (Figure 7D, *P*<0.01 for all the genes).

In addition to the regions detected by acH4 peaks, other genes were also over-expressed (~3400) or repressed (~2100) after NaBu treatment (see a list in **Table 4 and 5**). This effect confirms a major gene expression profile alteration after HDAC inhibition, being compatible with the enrichment obtained in the chIP-seq analysis for transcription factors. In this way, the changes in the histone H4 acetylation in several transcription factors would lead to an amplified gene expression response. Gene Ontology terms enriched either in the up-regulated or the down-regulated genes were unique, showing that major cellular pathways are altered in NaBu-treated animals. Biological processes that are up-regulated include protein phosphorylation and signaling and RNA metabolism, in additional to transcriptional regulation, which was already predicted using the chIP-seq data. Down-regulated processes include many metabolic pathways as well as cognition, response to light and muscle cell development, among others (Figure 7E and **F**, and **Table 6** and **7**).

In summary, we have shown that behavioral inter-individual variability depends on a regulatory pathway that affects histone H4 acetylation. In control conditions, YY1/HDAC1 complex deacetylates the histone H4 content of several regions located near genes coding for transcription factors. These changes, probably happening at the specific H4K12 mark, would lead to alterations in the gene expression profiles that might result into individual differences in their behavior. In the case of alteration of the YY1/HDAC1 pathway, these candidate regions are hyperacetylated and subsequently the neighboring genes overexpressed, leading to decreased inter-individual differences (Figure 7G).

## DISCUSSION

In this paper, we have found that a histone H4 acetylation pathway modulates individual behavior in a genetic-independent manner without affecting the global average behavior of the population. Histone H4 acetylation levels of an individual correlated with its individual behavior compared to the average of the population. Therefore, while the average behavior might depend more on genetic background (as seen for different strains in Figure 2) or environmental changes (as seen for different responses in Figure EV2D), behavioral inter-individual variability could result from histone H4 acetylation differences.

Several important questions arise from these results. The origin of genetic-independent changes in the individuals of an inbred population is still unknown. Our results suggest the existence of a stochastic basis for the generation of these individual differences. Several stochastic mechanisms could underlie behavioral inter-individual variability, such as paternal and maternal effects (Öst *et al*, 2014), differences in the experience received by individuals (Freund *et al*, 2013), or developmental noise, among others. Our results are consistent with a possible role of stochastic transcription factor binding to a set of target regions as a mechanism to generate transcriptional variability. Chromatin modifiers and histone marks have been shown to specifically affect gene expression noise (Weinberger *et al*, 2012; Wu *et al*, 2017). Thus, future studies will address the function of the YY1/HDAC1 complex in order to determine its binding dynamics. In addition, endogenous butyrate levels have been shown to be responsible of changes in the behavior within a microbiota-gut-brain link (Stilling *et al*, 2016). ß-hydroxybutyrate arising from lipid metabolism has been also found to endogenously inhibit HDACs, being the brain one of the main target tissues (Shirakawa *et al*, 2012). It will be interesting to study inter-individual differences in terms of microbiota and metabolomic content, and their possible relation to behavior.

Another open question is how histone H4 acetylation changes could lead to behavioral inter-individual variability. We found that histone H4 acetylation levels are functionally transformed into changes in gene expression. In addition, genes located near the candidate regions are significantly related to transcriptional regulation, so differences in their expression might be amplified and this ultimately could lead to differences in processes like cognition or visual response, as our RNA-seq results suggested. Previous work has described how inter-individual variability in other behavioral paradigms underlies in different physiological processes like neurogenesis (Freund *et al*, 2013; Simola *et al*, 2015) and serotonin signaling (Kain *et al*, 2012; Pantoja *et al*, 2016). As some of the target genes of the YY1/HDAC1 pathway are related to neurogenesis, like *neurog1* (Sun *et al*, 2001), it will be interesting to study if gene expression differences in these transcription factors could lead to alterations in neurogenesis and/or serotonin pathway. In addition, we have used activity and radial index as the parameters to quantify free-swimming behavior. As those might become proxy measures of anxiety levels and exploratory function in zebrafish (Kalueff *et al*, 2013), is even more relevant to characterize the serotonin pathway in the context of inter-individual variability in zebrafish.

Finally, we want to remark an interesting question that also arises from our data. A group of fish with high levels of histone H4 acetylation will behave more similarly one another than a different group of fish with lower levels of histone H4 acetylation. A potential explanation might be that the histone H4 acetylation (and consequently the gene expression) profiles become more different as their total levels decrease, due to stochastic binding to the target regions. Nevertheless, we cannot exclude that changes can occur at the cellular and/or the tissue levels, or that gene interactions participated in the generation of inter-individual differences. A future combination of experiments and theoretical modeling might clarify the generation of inter-individual variability by differences in histone H4 acetylation levels.

## METHODS

### Zebrafish lines and care

Zebrafish (*Danio rerio*) WIK strain (Nechiporuk *et al*, 1999) was kindly provided by Dr. Bovolenta (CBM-UAM) and inbred in our laboratory for at least three generations before the experiments. Afterwards, WIK F1 population was generated from a single batch of embryos from a single couple of adult fish. Two additional cycles of inbreeding were carried out, crossing a couple of siblings from the former generation. CG2 (Mizgirev & Revskoy, 2010) clone population, generated by double gymnogenetic heat-shock, and characterized by being pure isogenic zebrafish was kindly provided by Dr. Revskoy (Univ Northwestern) as a control of reduced genetic differences between siblings. The outbred LPS (Local Pet Store) strain was recently described (Pérez-Escudero *et al*, 2014), and used as a model of genetic heterogeneity. Heterozygotic *hdac1 (hi1618)* and *yy1a (sa7439)* mutant strains with wild-type (AB strain) counterparts were obtained from ZIRC, while heterozygotic *hdac1* (*sa436*) (Noël *et al*, 2008) with wild-type counterparts were kindly provided by Dr. Ober (University of Copenhagen).

Care and breeding of the zebrafish strains were as described (Pérez-Escudero *et al*, 2014), with specific additional details. Eggs were isolated after 24 hours post-fertilization, and maintained in custom multiwell plates until 10 days post-fertilization (dpf). They were fed (JBL NovoBaby) from 6 dpf and water was changed daily if it is not indicated specifically in the experiment.

All the experiments using animals were approved and performed following the guidelines of the CSIC (Spain) and the Fundaçao Champalimaud (Portugal) for animal bioethics.

### Free-swimming setup and recording

The setup consists of a monochrome camera located over the wells at a distance of 70 cm and pointing downwards. The camera used was a 1.2 MPixel camera (Basler A622f, with a Pentax objective of focal length 16 mm). The wells are circular, carved on transparent PMMA (24 wells per plate, and typically two plates are recorded simultaneously), and have their walls tilted so that even in the most lateral wells the wall never hides the larva from the camera. Each well is 15 mm deep, and has a diameter of 1.8 mm at the bottom and a diameter of 30 mm at the top (**Figure S1A**). For the experiments, each well is filled with a volume of 3 ml. The dishes are supported by a white PMMA surface that is only partially opaque. Behind this white surface we place two infrared led arrays (830nm, TSHG8400 Vishay Semiconductors) pointing outwards (Figure EV1A). Two paper sheets stand between the lights and the central space that lies directly under the wells. With this disposition we ensure that only diffuse indirect light reaches the wells, so that the illumination is roughly uniform (most of the light comes from below the wells through the white surface). All the set-up is surrounded by white curtains. Video camera recorded at a 25 fps rate (Figures EV1B, EV1C for examples of a single frame and final trajectories).

A larval population (5-8 dpf) consisted of at least 24 fish siblings from the same batch of embryos. After five minutes of acclimation to the new environment, the larvae were recorded for 20 minutes. Water temperature was maintained in a strict range (27-28 ºC) during each experiment.

### Custom-built software tracking larvae

We developed multiwellTracker, a software to automatically track zebrafish larva in wells. The software is available at http://www.multiwelltracker.es.

#### Detection of wells

The program is prepared to auto-detect circular wells, regardless of their spatial arrangement. To detect the wells we use the circular Hough transform (we have modified the code of Tao Peng distributed by Matlab Central under BSD license). In order to estimate the diameter of the wells, it computes the image’s Hough transform for 100 radii different in 5 pixels and a rough estimate of the largest possible radius (length of the longer side of the image divided by the square root of the number of wells) (Figure EV1D). The system selects the highest point of this measure as an estimate of the radius of the wells (r_est_). It is possible to skip this first step and instead specify manually a value for r_est_. This may be advisable when many videos are recorded with the same set-up and the same wells.

In the second step the system locates the centers of the wells. To do this it performs a Hough transform of the original image, this time with radii only in the range between 0.8r_est_ and 1.2r_est_. The transformed image usually has clear peaks in the centers of the wells. Then it filters the transformed image with a Gaussian filter to increase its smoothness (the resulting transformed image is shown in Figure EV1E). Then, it selects the maximum of the transformed image as the center of the first well. To prevent selecting the same well twice, the system discards all the pixels of the transformed image that are within radius r_est_ of the selected center (Figure EV1F). It selects the new maximum as the center of the second well, and repeats the procedure until all wells have been found (Figure EV1G). The experimenter can correct the result by manually clicking on the center of the wells that have not been correctly located (<1% of cases).

#### Pre-processing of images

In order to control for fluctuations in illumination, each frame is normalized by dividing the intensity of each pixel by the average intensity across all pixels of the frame. After normalizing the frame, a 2D Gaussian filter is used to smooth the image (Figure EV1H shows the image before and Figure EV1I after filtering).

#### Background subtraction and detection of the larva

In order to extract the image of the larva from the background, the system finds the average of 1,000 frames equi-spaced along the whole video. This average image is what we will call “static background”. By subtracting each frame by the static background, we obtain an image in which the larvae correspond to dark regions (Figure EV1J). However, because of relatively slow changes in the set-up over time, the system uses the static background in combination with a dynamic background, which is computed as the average of the previous 5 frames. The difference between the current frame and the dynamic background will only show larvae that are moving in that precise moment (Figure EV1K).

The specific algorithm to detect the larva is as follows. First, the difference between the current frame and the static background is thresholded keeping only pixels for which the difference is below −0.5. We then find connected components (“blobs”) in this thresholded image, keeping those that are larger than 1 pixel. Because these blobs come from the difference with the static background, both static and moving larvae will be detected. But at this stage some blobs come simply from noise. In order to filter out noisy blobs, the system accepts a blob if it fulfills at least one of these two conditions: (a) It contains at least one pixel that was identified as part of the larva in the previous frame or (b) it contains at least one pixel for which the difference between the current frame minus the dynamic background is below the same threshold as before (−0.5).

#### Removal of reflections

In most cases only one blob is obtained after the process described in previous sections. But when the larva is close to the wall of the well, its reflection on the wall may also be selected. The system considers that a blob A is a reflection of blob B when all of the following conditions are met: (a) Blob B is bigger than blob A, (b) blob B is closer to the center of the well than blob A and (c) the lines between the center of the well and the two blobs form an angle < 10°. When these three conditions are met, the system removes blob A.

#### Acquisition of the position of the larvae

If more than one blob remains in the same well after the previous steps, the system selects the one with highest contrast. For the selected blob, the system takes the position of its most contrasted pixel, and adds this position to the trajectory of the larva. If in a well no blob remains after the previous steps, the trajectory is left with a gap. When this happens, the program will not re-track the larva until it moves.

### Behavioral Parameters

Different parameters reflecting the behavior of individual larvae were measured, and finally two of them were used through the paper: (i) activity (percentage of time in movement) and (ii) radial index (average position from the border towards the center of the well). We also studied three additional parameters: (i) Tortuosity in the trajectory was calculated as the scalar product of the velocity vectors between two consecutive frames and the value in Figure EV1L was obtained by averaging this parameter through the whole video, excluding the frames where the animal was immobile. (ii) Speed was calculated as the average distance (in pixels) travelled per frame, in those frames where the fish was active. (iii) Bursting was obtained as the total number of frames where fish changed from immobility to motion. We found that these three parameters correlated with activity.

The average of each individual parameter was tested from 5 to 8 days post-fertilization (dpf) to assess if individual behavior was significantly stable along the days using Pearson coefficient of correlation.

### Additional validation of the experimental setup

Several controls were performed for possible experimental artifacts affecting wells differently. Behavioral parameters were robust to 90 degrees counterclockwise rotations of the multi-well plate (Figure EV2A, left, R=0.73, *P*<0.001, and R=0.68, *P*<0.001; for linear correlation tests of individual behavior) or to interchanging the larvae between outer and inner wells (Figure EV2A, right, R=0.65, *P*<0.001, and R=0.61, *P*<0.001; using the same correlation test). Also, we found no correlation between the small differences in illumination across wells and behavior (Figure EV2B). We further corroborated using a significance test that the differences in behavior did not have an origin in systematic differences across wells. For this, we found that the average behavioral parameters obtained in fifteen individual experiments were not different between wells (Figure EV2C, typically *P*=0.4 and always *P*>0.19 for both parameters).

### Stimulus response tests

We studied the influence that our free-swimming behavioral parameters could have on the performance of the individuals when they respond to three different stimuli.

#### Response to mechanical disturbance

We applied mechanical perturbations to each larva by pipetting up and down the water content of the well for four times. Perturbations were applied at 6 dpf to previously recorded animals, and the 20-minute recording was done at 7 dpf. The recording was performed in the usual setup.

#### Response to strong Light Pulse

In complete darkness, we applied three different light flashes to the larvae and study their behavior in the 90 subsequent seconds. The flashes and the recording were performed in the usual setup. Pre-recording behavioral parameters were obtained the day before.

#### Novel tank with light/dark preference

In order to study the effect that a novel setup could have on the behavior of larvae we built a rectangular setup, which changed the geometry of the previous circular wells. The setup dimensions were 84 mm x 21 mm and it was built in transparent acrylic. To try to see if our parameters had any effect on the light-dark preference, half of the floor of the setup (42 mm x 21 mm) was white while the other half was black. The height of the setup was 5 mm. Larvae were placed in the center of the white part and recorded for 10 minutes. Activity was calculated as previously described and distance to the wall was represented by the average distance to the longest walls, normalized to 1 in the middle point of both walls and to 0 in the exact position of the walls.

The effect of our behavioral tests resulted in a decrease (increase) in mean activity (radial index), but maintaining the same individuality of the pre-recorded free-swimming experiments (Figure EV2D; *P*<0.04 for changes in mean activity and radial index compared to control larvae of the same age; *P*<0.02 for linear correlation of activity and radial index. In the case of novel tank, radial index cannot be applied because the wells are elongated and was then replaced by the minimum distance to the longer walls. We note, however, that this parameter showed no correlation with the radial index of pre-recorded experiments in the same animals.

### Inter-individual vs. Intra-individual differences

The behavioral parameters (activity and radial index) were also obtained from consecutive fragments of 30 seconds for each 20-minute experiment for each larva. This was fitted to a two-dimensional Gaussian, but for clarity when representing many animals (like in Figure 1E) an isocontour of the Gaussian for each animal was used. An isocontour is an ellipse with principal axes given by the eigenvectors of the covariance matrix. We chose the isocontour with length of each semiaxis given by the square root of the eigenvalue of the covariance matrix, as this reduces to the standard deviation in each direction for cases with no correlation between the two variables. Intra-individual variation distribution was obtained using the coefficients of variations (CVs) for activity and radial index separately. Inter-individual variation was calculated the same way but using fragments from different fish.

### Comparing the behavioral variability between two animal groups

A simple visual method to characterize the variability in a population is to plot the bi-dimensional distribution of the activity and radial index of individuals (like in Figure 1F). To do so, we used Gaussian kernel smoothing that consists in adding up Gaussians centered at the data points as

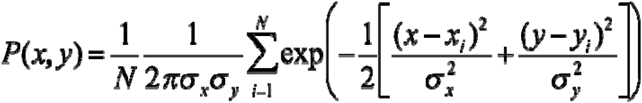

with *x*_i_ and *y*_i_ the mean activity and radial index values of individual *i* of a total of *N* members of the population. An optimal smoothing uses standard deviations of each Gaussian given by **σ − *N*^−1/6^α_*x*_** with **α_*x*_** the standard deviation in the *x*_i_ data values, and similarly for **σ_*y*_** using the *y*_i_ values (see B.E. Hansen, unpublished manuscript, http://www.ssc.wisc.edu/~bhansen/718/NonParametrics1.pdf). The volume under the probability surface has a value of 1, even when the values of the probability density are already up to 90. The probability surface sits on an area on the x-y plane of approximately 0.4x0.4, making the total volume under the surface to be 1.

While this distribution gives a visual and intuitive characterization of behavioral variability, an even simpler characterization is achieved using, for each group, a single parameter summarizing its two-dimensional variability. We used *generalized variance* as this single parameter, measured as the determinant of the covariance matrix (**Table 1**), while other parameters like the standard deviation for each parameter gave similar statistical results (**Table 2**).

### ChIP-seq, conventional chIP, and rechIP analyses

Chromatin immunoprecipitation was obtained from pooled samples of at least four zebrafish larvae. Briefly, the samples were crosslinked with 1.8% formaldehyde for 30’ and then quenched with 1% glycine for 5’. Extracts were lysed using a SDS Lysis buffer (50 mM Tris-HCl pH 8.1, 1% SDS, 10 mM EDTA) for 30’ at 4ºC, and then diluted with a Dilution buffer (6.7 mM Tris-HCl pH 8.1, 0.01% SDS, 1.2 mM EDTA, 1.1% Triton X-100, 167 mM NaCl). 2 mM sodium butyrate was added to avoid histone deacetylation activity during the preparation. Then, the fish were sonicated with two pulses (30’’ ON / 30’’ OFF) of 15’ each with the Diagenode Bioruptor. Before pre-clearing the samples with protein A/G beads, an input sample was obtained. Then, the extracts were immunoprecipitated overnight using 1 μg of the anti-acetyl-Histone 4, anti-HDAC1 or anti-YY1 antibodies. Bound DNA was recovered with protein A/G beads, then washed with Low-Salt (120 mM Tris-HCl pH 8, 0.1% SDS, 2 mM EDTA, 1% Triton X-100, 150 mM NaCl), High-Salt (120 mM Tris-HCl pH8, 0.1% SDS, 2 mM EDTA, 1% Triton X-100, 500 mM NaCl), LiCl (10 mM Tris-HCl pH 10, 1 mM EDTA, 0.25 M LiCl, 1% NP40, 1% sodium deoxycholate) and two times with 1X TE (10 mM Tris-HCl pH 8, 1 mM EDTA) buffers, and recovered with Elution (1% SDS, 0.1 M NaHCO3). DNA purified samples were de-crosslinked using sodium chloride, and cleared with Qiagen spin columns, and for rechIP, the samples were reincubated with a second antibody after different elution (Román *et al*, 2011), and another round of washes and elution was performed.

In the case of conventional chIP, qPCR was used to detect differences between the samples in the target regions, using the unbound fraction (chIP from the same sample but without antibody) as a control to normalize results.

In the case of the acetyl-H4 chIP-seq, we prepared twelve samples (4 control, 4 NaBu and 4 random) composed of four fish each. The larvae selected for each sample were chosen by the following algorithm. In the first (control) cluster, a Hierarchical Clustering analysis using Euclidean distance as the metric and the average linkage clustering as the linkage criteria was used the clustering in the chIP-seq, as the total population consisted of 72 larvae. The selection in the NaBu experiment used the same algorithm. In the random experiment, fish were randomly selected from the population.

The final samples were processed at the Genomics Unit at the Scientific Park of Madrid. Libraries were built, and the samples were sequenced using an Illumina GAIIX. Reads were aligned to *Danio rerio* genome sequence (Zv9) with BWA, and the results were subjected to the MACS peak detection algorithm (Feng *et al*, 2012). Afterwards, peaks from the different samples were merged and quantified separately as Fragments Per Kilobase per Million reads (FPKM) using DiffBind package (Caryn *et al*, 2012), obtaining 27,310 peaks in Control samples and 33,649 peaks in NaBu samples, with 12,419 peaks being shared by both conditions. Finally, EdgeR (Robinson *et al*, 2010) was applied to detect differential binding between control and NaBu samples. The candidate peaks with higher histone H4 acetylation in NaBu compared to control (*P*<0.001) were further filtered using the random samples. Specifically, we removed the regions with higher variability (measured with the Coefficient of Variation) in random samples, as this variability is not associated to behavior.

### Gene Ontology and Transcription Factor Binding Sites analysis

Position of the candidate peaks obtained in the chIP-seq was retrieved in order to study their relative position to near genes. Nearest genes were retrieved, and they were analyzed for Gene Ontology using DAVID (Dennis *et al*, 2003) as in the case of RNA-seq targets. In addition, the candidate regions located near the TSS of a gene were used to predict enriched DNA motifs and their potential biological activity with MEME suite (Bailey *et al*, 2009).

### Reagents and antibodies

Sodium butyrate (NaBu), trichostatin A (TSA), cambinol (Cmb) and 5-Aza-2′-deoxycytidine (AZA) were purchased from Sigma-Aldrich (#303410, T8552, C0494 and A3656), and then they were dissolved in Phosphate-buffered saline (PBS) to be used in a final 2 mM, 0.1 µM, 0.2 µM and 15 mM concentration of fish water, respectively. PBS alone was used as vehicle control. The pharmacological treatment lasted for 24 hours from 7 dpf to 8dpf. Acetyl-Histone 4 and acetyl-Lysine antibodies were obtained from Millipore (#06-866 and #05-515), anti-HDAC1 and anti-YY1 from ActiveMotif (#39531 and #61780), anti-Sp1 and anti-H4 from Abcam (ab59257 and ab16483), and McrBC enzyme from New England Biolabs (M0272).

### Immunoprecipitation and Western Immunoblotting

Groups of twenty fish (control, NaBu-treated, *hdac1* +/− and *yy1* +/−) were frozen at 8 dpf. They were lysed in a solution containing 100 mM TrisHCl pH 7.5, 20 mM NaF, 2 mM DTT, 2 mM EGTA, 2 mM EDTA, 1 mM sodium orthovanadate, 0,54M sucrose, 0.2 mM phenyl-methyl sulfonyl fluoride, 2% X-100 Triton, ß-Mercaptoethanol and 4 μg/ml Complete protease inhibitor cocktail (Roche, #11836153001). For each condition/treatment an aliquot of 1 mg protein was incubated with 2 μg of anti-YY1 antibody and 30 μl of protein-A/G plus Sepharose beads overnight at 4°C. Beads were then washed five times with washing buffer (20 mM Tris–HCl pH 7.4, 50 mM NaCl and 4 μg/ml Complete protease inhibitor cocktail). Immunoprecipitated proteins were analyzed by western-blot using anti-acetylated lysine antibody and YY1 antibody. Immunoprecipitated proteins were analyzed by western-blot using anti-acetylated lysine antibody and YY1 antibody.

### RNA isolation, qPCR quantification and RNA-seq

Total RNA was isolated using homogenized extracts from at least five fish per sample by RNeasy Mini (Qiagen, #74104) purification. Retrotranscription was done with iScript (Bio-Rad, #1708891) following manufacturer recommendations. Finally, quantification of the target genes was measured using qPCR with specific primers. Values were normalized using the GAPDH results obtained from the same sample, and *P*-values obtained by using Student’s T-test. In the case of RNA-seq, samples were processed at the Genomics Unit at the Scientific Park of Madrid to generate libraries, and raw reads were obtained using Illumina GAII. Afterwards, the reads were aligned to *Danio rerio* genome (Zv9) using BWA, and transcript counts were normalized to TMM (Trimmed Mean of M-values) scores. Then NoiseqBIO algorithm from Noiseq was used to detect significantly (adjusted P-value<0.1) altered genes in the two groups.

### Quantification of histone acetylation levels

Eighteen clusters of five fish from a total population of 90 were obtained from the behavioral space (activity/radial index) using an *ad hoc* algorithm. First, 18 centroids were randomly chosen, and 5 individuals were assigned to the nearest (not occupied) centroid. Then, centroids were redefined using the average values of the new clusters, and a new round of assignment of the fish to the centroids was done. This iteration was repeated until the centroids were stable. Then, total histones were extracted using an Epigentek kit (OP-0006) and quantified by Pierce Coomasie Plus reagent (Thermofisher, #27236), in order to use the same amount of total histones in each sample. Acetyl-H4, acetyl-H3 and specific acH4 marks (H4K5, H4K8, H4K12 and H4K16) were quantified by ELISA following manufacturer recommendations using the Epigentek kits P-4009, P-4008 and P-3102, respectively.

### Quantification of methylated DNA

DNA methylation was quantified using larval DNA digested by MCrBC enzyme as previously done (Torres-Núñez *et al*, 2015) following kit instructions.

### Statistical analysis and data storage

In the case of behavioral parameters and bulk acetylated histone measures, the statistical tests to compare the differences between two distributions were conducted by calculating the value of their representative parameter (*i.e*., generalized variance, Pearson’s correlation coefficient…), shuffling randomly the data of both distributions for 1,000 – 10,000 times and computing a *P*-value given by the proportion of times in which the difference in the representative parameter of the random distributions was higher than their original value.

In the case of chIP, rechIP and RNA analyses, the statistical tests to compare the difference between two conditions were conducted by calculating the representative parameter (fold change compared to a control), and *P-*values were obtained using the Student’s T-test.

All the experiments (except chIP-seq and RNA-seq, in which a FDR<=0.05 was applied) were done at least three times with different biological datasets, and *P*-values were calculated using the three replicas. Figures show a representative experiment of the triplicate. MATLAB and R were used for all the computations and the statistical analysis.

ChIP-seq and RNA-seq data are stored in ArrayExpress repository (E-MTAB-5993 and E-MTAB-5992, respectively). All the video trajectories of the fish experiments presented in the manuscript are available upon request.

## AUTHOR CONTRIBUTIONS

A.-C.R. designed project, performed experiments and analysis and wrote the paper, J.V.-P. performed experiments and analysis and wrote the paper, A.P.-E. developed the tracking software, J.M.C.-G. performed experiments, P.M.F.S provided reagents, and G.G.d.P. designed and supervised project, performed analysis and wrote the paper.

## ACKNOWLEDGEMENTS

We thank Rui Costa, Carlos Ribeiro, Antonia Groneberg, and other members of the Champalimaud Neuroscience Programme for discussions, and Isidro Dompablo (CSIC) and Ana Catarina Certal (Champalimaud Foundation) for technical assistance and animal care. CG2, *hdac1 sa436*, and WIK lines were kind gifts from Sergei Revskoy, Elke Ober and Paola Bovolenta, respectively. This work was supported by a Juan de la Cierva and a FEBS Long-Term fellowship (to A.-C.R.), a JAE-CSIC Predoctoral Fellowship (to J.V.P.), Spanish Plan Nacional Ministerio de Economia y Competitividad grants BFU2014-54699-P (to J.M.C.-G.), BFU2011-22678 (to P.M.F.S), BFU2012-33448 (to G.G.d.P.), ERASysBio+ initiative supported under the European Union European Research Area Networks (ERA-NET) Plus scheme in Framework Program 7 (to G.G.d.P.), Fundaçao para a Ciência e Tecnologia PTDC/NEUSCC/0948/2014 (to G.G.d.P.) and Champalimaud Foundation (to G.G.d.P.). The funders had no role in study design, data collection and analysis, decision to publish, or preparation of the manuscript.

## EXPANDED VIEW FIGURE LEGENDS

**Figure EV1.**
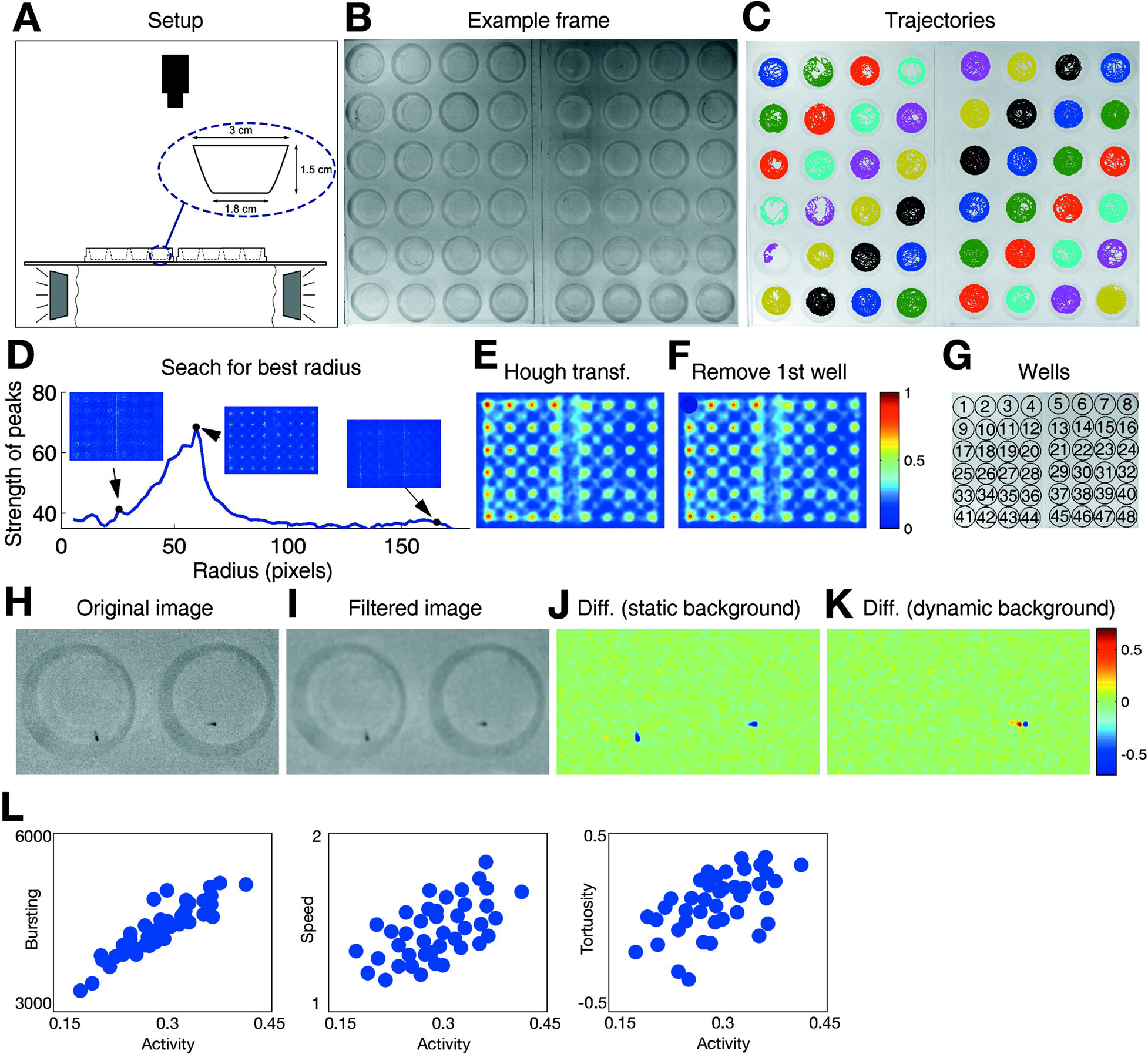
Setup, tracking system and behavioral parameters. **(A)** Scheme of the setup with an enlarged scheme of one of the wells. **(B)** Image of the wells. **(C)** Trajectory of each larva. **(D)** Strength of the peaks resulting from Hough transforms with different radii. The insets show the Hough transforms for 3 different radii. From this calculation we define rest = 58 pixels. **(E)** Hough transform of the frame, using radii in a small interval around rest (in this case, in the interval between 47 and 71 pixels). **(F)** Same as **E**, but after removing the region that corresponds to the highest maximum. **(G)** Final result of the detection of wells. **(H)** Original image of two adjacent wells. **(I)** Image of the same two wells, after filtering with a Gaussian filter. **(J)** Difference between the filtered image of the two wells, and the static background. The two larvae are clearly visible as regions with low values. **(K)** Difference between the filtered image of the two wells, and the dynamic background. Only the larva on the right is visible, because the larva on the left is not moving in this specific frame. **(L)** Correlation between the different behavioral parameters analyzed (Bursting, Speed and Tortuosity) and activity.

**Figure EV2.**
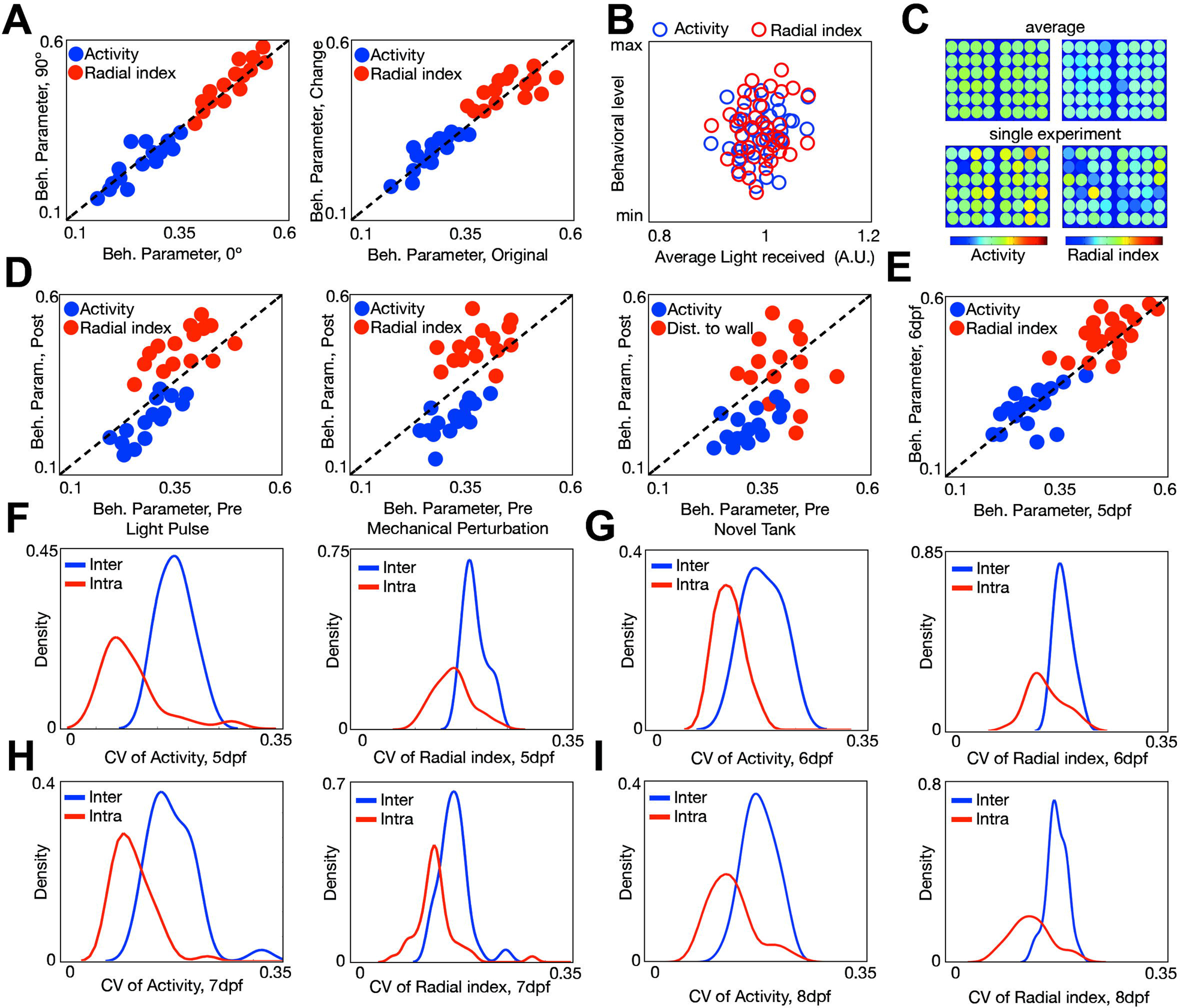
Additional behavioral parameters for validation of the setup. **(A)** Changes in the behavior after 90º rotation of the plates (left) and after interchanging 24 larvae from external to internal wells (right). **(B)** Larval behavioral parameters (activity, blue; radial index, red) plotted against the average light received by these 48 larvae. Min and max values are 0.1−0.5 and 0.3−0.7 for activity and radial index, respectively. **(C)** Average activity (left) and radial index (right) of a pool of different larvae recorded in the same wells for 15 experiments (>600 larvae) (top), compared to a single 48 larval population (bottom). **(D)** Correlation between the behavior in different tests (Light Pulse, left; Mechanical Perturbation, center and Novel Tank with Light Preference, right) and our behavioral parameters in the usual conditions, using 16 larvae. **(E)** Correlation of activity (blue) and radial index (red) between 5 dpf and 6 dpf for the same group as in Figure 1. **(F)** Smoothed histogram of the coefficient of variation of the activity (left) and the radial index (right) showing the intra-individual variability (red) and inter-individual variability (blue) of the same 48 larval group shown in Figure 1 at 5 dpf. **(G)** Same as **F**, for the group at 6 dpf. **(H)** Same as **F**, for the group at 7 dpf. **(I)** Same as **F**, for the group at 8 dpf.

**Figure EV3.**
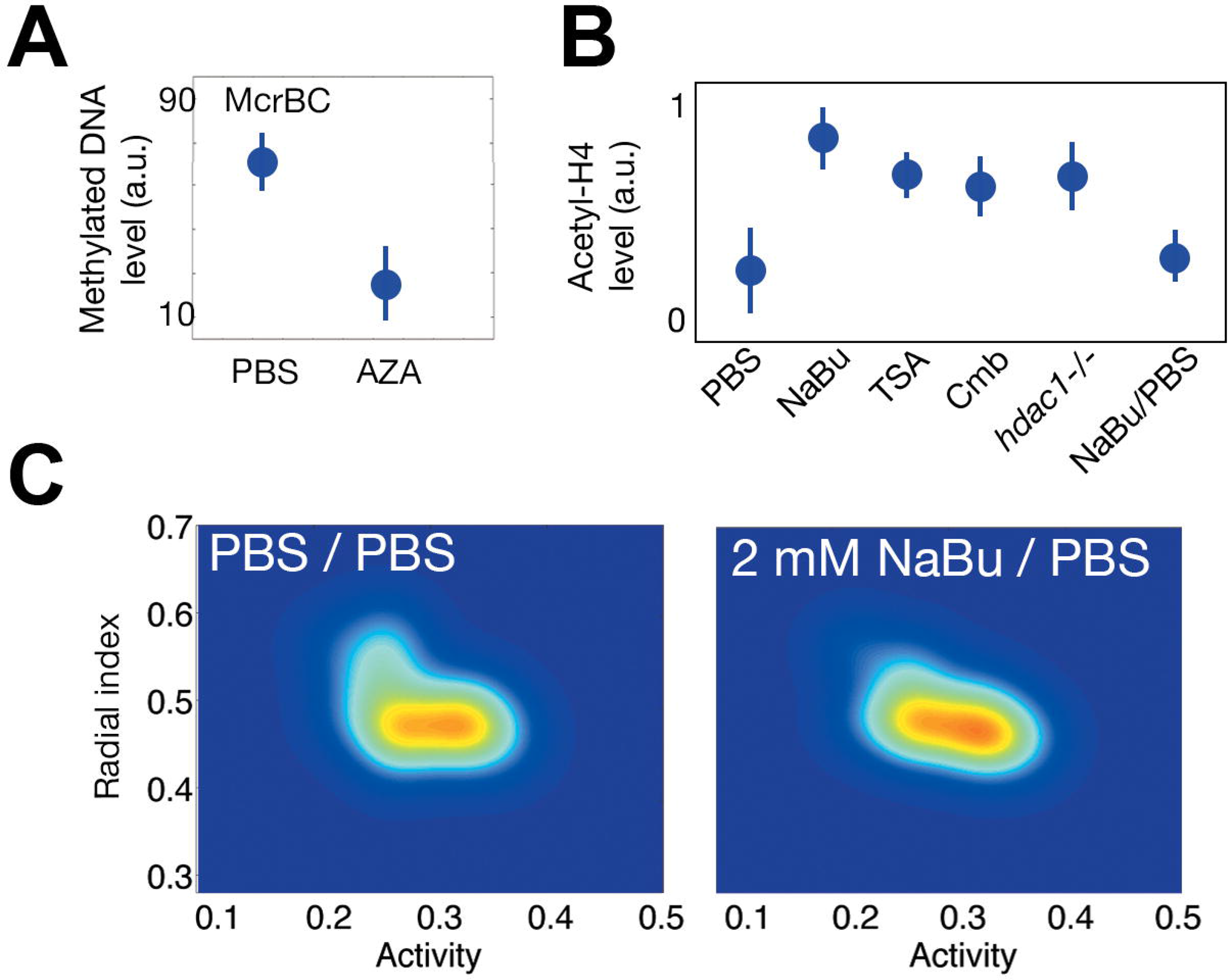
Additional data for epigenetic treatments. **(A)** Methylation levels comparing control (PBS) and AZA treatment. **(B)** ELISA results quantifying the levels of acetyl-H4 in the conditions shown in Figure 3. **(C)** Probability density for 24 larvae 48h after treatment with NaBu only during the first 24h and then washed with PBS (PBS/PBS as control).

**Figure EV4.**
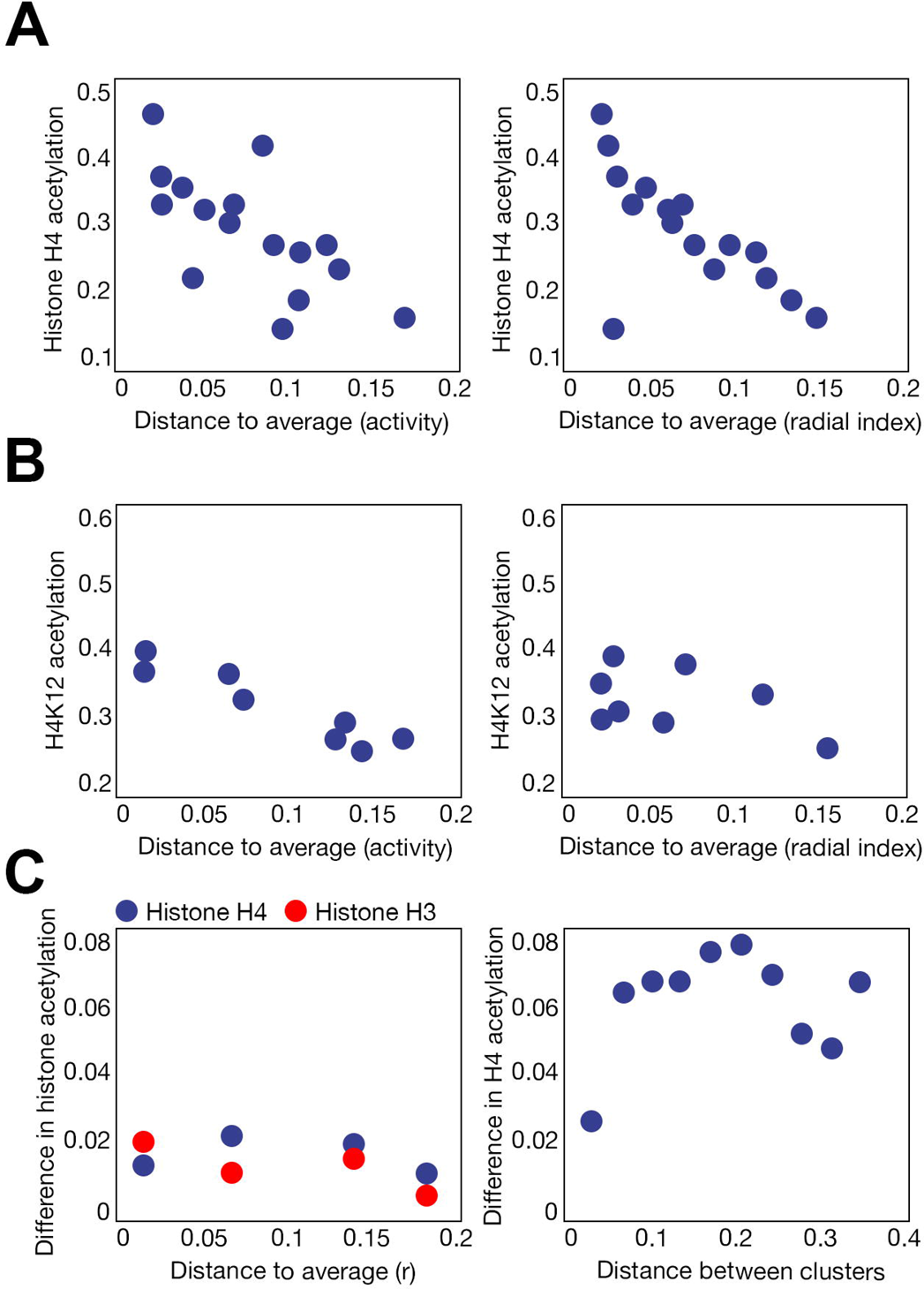
Additional data for histone H4 acetylation and behavior. **(A)** Relation between histone H4 acetylation levels and distance to average behavior: activity (left) and radial index (right). (**B**) Same as **A**, but for H4K12 acetylation levels. (**C**) Differences in histone H3 (red) and H4 (blue) acetylation levels within extracts obtained in the same behavioral positions towards the average behavior of the population (left). Difference in histone H4 acetylation in clusters with different behavioral positions towards the average behavior of the population (right).

**Figure EV5.**
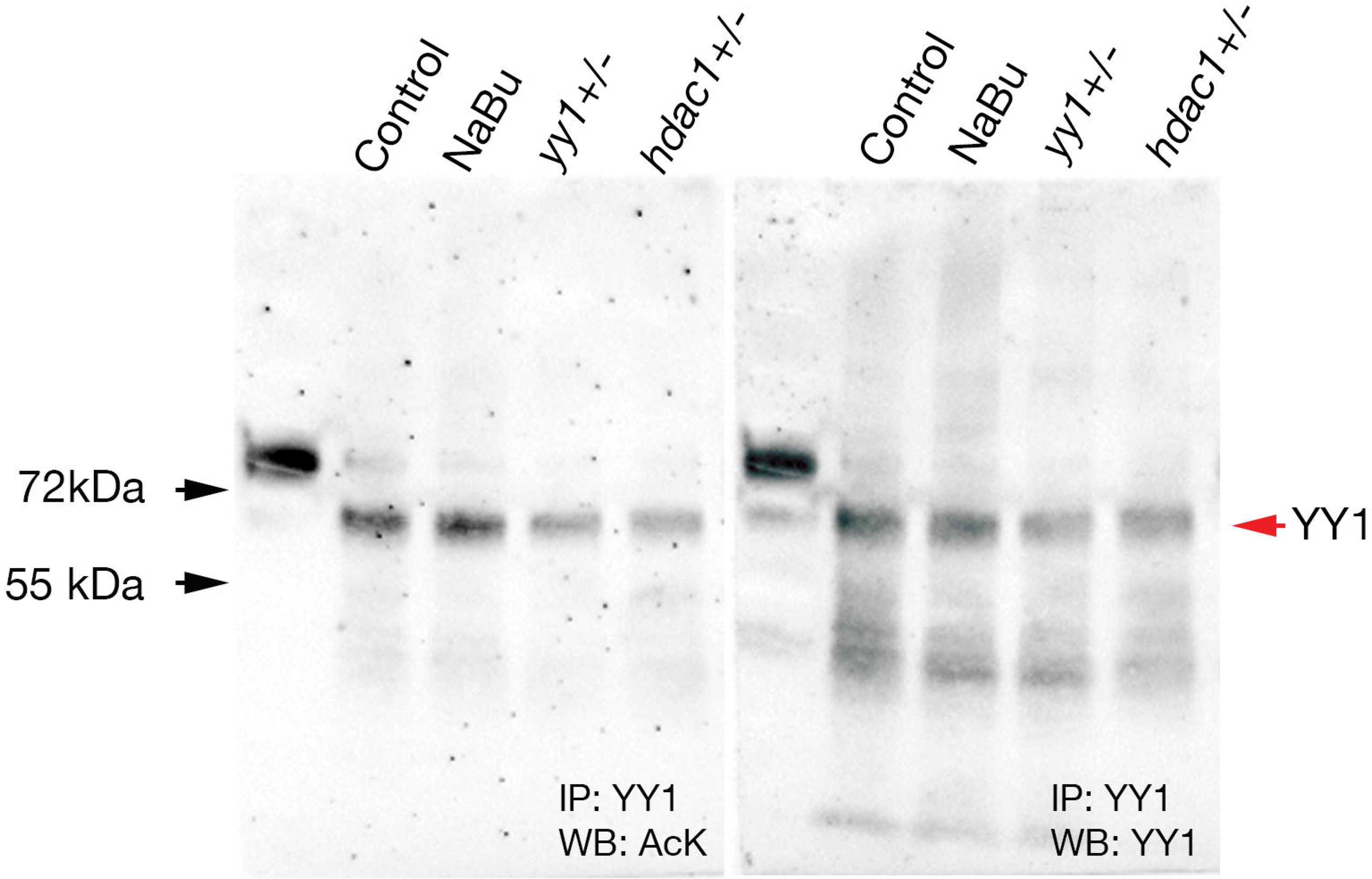
Additional data for acetyl-YY1 experiment. Original blot from the figure 6J representing the acetyl-YY1 levels at different conditions.

